# Rapid Quantification of Peptide Oxidation Isomers from Complex Mixtures

**DOI:** 10.1101/850032

**Authors:** Niloofar Abolhasani Khaje, Joshua S. Sharp

**Affiliations:** Department of BioMolecular Sciences, School of Pharmacy, University of Mississippi, University, Mississippi 38677; Depertmant of Chemistry and Biochemistry, University of Mississippi, University, Mississippi 38677

## Abstract

Hydroxyl radical protein footprinting (HRPF) is a powerful technique for probing changes in protein topography, based on quantifying the amount of oxidation of different regions of a protein. While quantification of HRPF oxidation at the peptide level is relatively common, quantification at the residue level is challenging due to the influence of oxidation on MS/MS fragmentation and the large number of complex and only partially chromatographically resolved isomeric peptide oxidation products. HRPF quantification of isomeric peptide oxidation products (where the peptide sequence is the same but isomeric oxidation products are formed at different sites) at the residue level by electron transfer dissociation tandem mass spectrometry (ETD MS/MS) has been demonstrated in both model peptides and HRPF products, but the method is hampered by the partial separation of oxidation isomers by reversed phase chromatography. This requires custom MS/MS methods to equally sample all isomeric oxidation products across their elution window, greatly increasing method development time and reducing the oxidation products quantified in a single LC-MS/MS run. Here we present a zwitterionic hydrophilic interaction capillary chromatography (ZIC-HILIC) method to ideally co-elute all isomeric peptide oxidation products while separating different peptides. This allows us to relatively quantify peptide oxidation isomers using an ETD MS/MS spectrum acquired at any point across the single peptide oxidation isomer peak, greatly simplifying data acquisition and data analysis at both the peptide and amino acid level.

## Introduction

Hydroxyl radical protein footprinting (HRPF) by fast photochemical oxidation of protein (FPOP)^1^ is a powerful technique to study changes in protein topography^2,3^. In FPOP, hydroxyl radicals react covalently with solvent exposed amino acid side chains at a rate influenced in part by the side chain’s solvent accessible surface area and the intrinsic reactivity of the amino acids themselves^3,4^. This reaction is an oxidative process, most commonly leading to the net addition of oxygen to amino acids, which is measured by liquid chromatography coupled to tandem mass spectrometry (LC-MS/MS)^5–10^. Fast generation of hydroxyl radicals by FPOP^11,12^ allowed benchtop HRPF to be successfully applied to a variety of applications including studying protein structure and conformational changes^3,13–18,^ protein-ligand interactions ^19–21^, protein-protein interactions^22,23^ and even in-cell and *in vivo* protein conformation^24,25^. This technology measures changes in the apparent rate of oxidation of various regions of the protein, and correlates those changes with changes in that region’s average solvent accessible surface area^5,26^. For example, in a protein ligand interaction study, as a ligand binds to a region of a protein and shields that region from solvent, the apparent rate of oxidation of that region measured in an FPOP experiment will decrease relative to the average amount of shielding. The spatial resolution of the technique is limited by the size of the region that can be accurately probed (e.g. a tryptic peptide). Despite all the advancement in HRPF development, this technique hasn’t been easily transferable to the broader community due in part to the complexity of data analysis and data acquisition. One of the reason for complexity of the data is because hydroxyl radicals often generate isomeric peptide oxidation products simultaneously^1,27,28^. Quantification of a mixture of isomeric peptide oxidation products (where the primary sequence is the same and the oxidation product has the exact same mass, but the modification occurs on different amino acids in that sequence) is challenging due to the influence of the modifications in ergodic fragmentation mechanisms^21,29,^ and the ability of chromatography to partially (but not fully) separate some peptide modification isomers^2,21,30,31^. Previous studies using synthetic isomeric peptide oxidation products and C18 reversed phase chromatography showed poor quantification by chromatographic peak area even when these isomers could be separated, probably due to non-ideal peak shapes from partially resolved sub-residue peptide oxidation isomers (including stereoisomeric oxidation products)^32^.

Recently, C18 reversed phase chromatography followed by electron transfer dissociation (ETD) MS/MS has been used to quantify modifications on each residue of a peptide for complex mixtures of peptide oxidation isomers in a technique termed high resolution hydroxyl radical protein footprinting (HR-HRPF)^21^. While ETD-based quantification remains the only method for oxidation isomer quantification by LC-MS that has been validated using defined mixtures of isomeric peptide oxidation products^32^, the data acquisition and quantification are still difficult due to the high-resolution hydrophobic separation of the peptides by reversed phase liquid chromatography (RPLC), resulting in peaks that do not co-elute and are therefore not sampled equally during data-dependent fragmentation^33^. In order to overcome C18 RPLC problems for ETD based fragmentation, a custom method has to be made for each experiment to ensure equal sampling during ETD fragmentation throughout the entire elution window of each modified peptide. Each sample has to be run first using data dependent acquisition method followed by manual determination of the elution window of each peptide and all of their oxidation products. ETD fragmentation must then be scheduled across the entire elution window of each set of peptide oxidation products, with fragmentation scan frequency sufficiently high to accurately sample the peak shape of each partially separated isomer. An example of a method to address the oxidation products of a single peptide is shown in Figure 1; often, it is necessary to make multiple injections to ensure ETD coverage of all oxidation isomers in even modestly complex samples. Making each custom method requires additional sample injections, analysis time, and time to create a custom method for each experiment. While the quantitative results provide accurate structural information^3,21^, it is also labor intensive, often requires multiple injection of sample to fully cover all peptide oxidation products, and is difficult to automate^3,21,23^.

**Figure 1.**
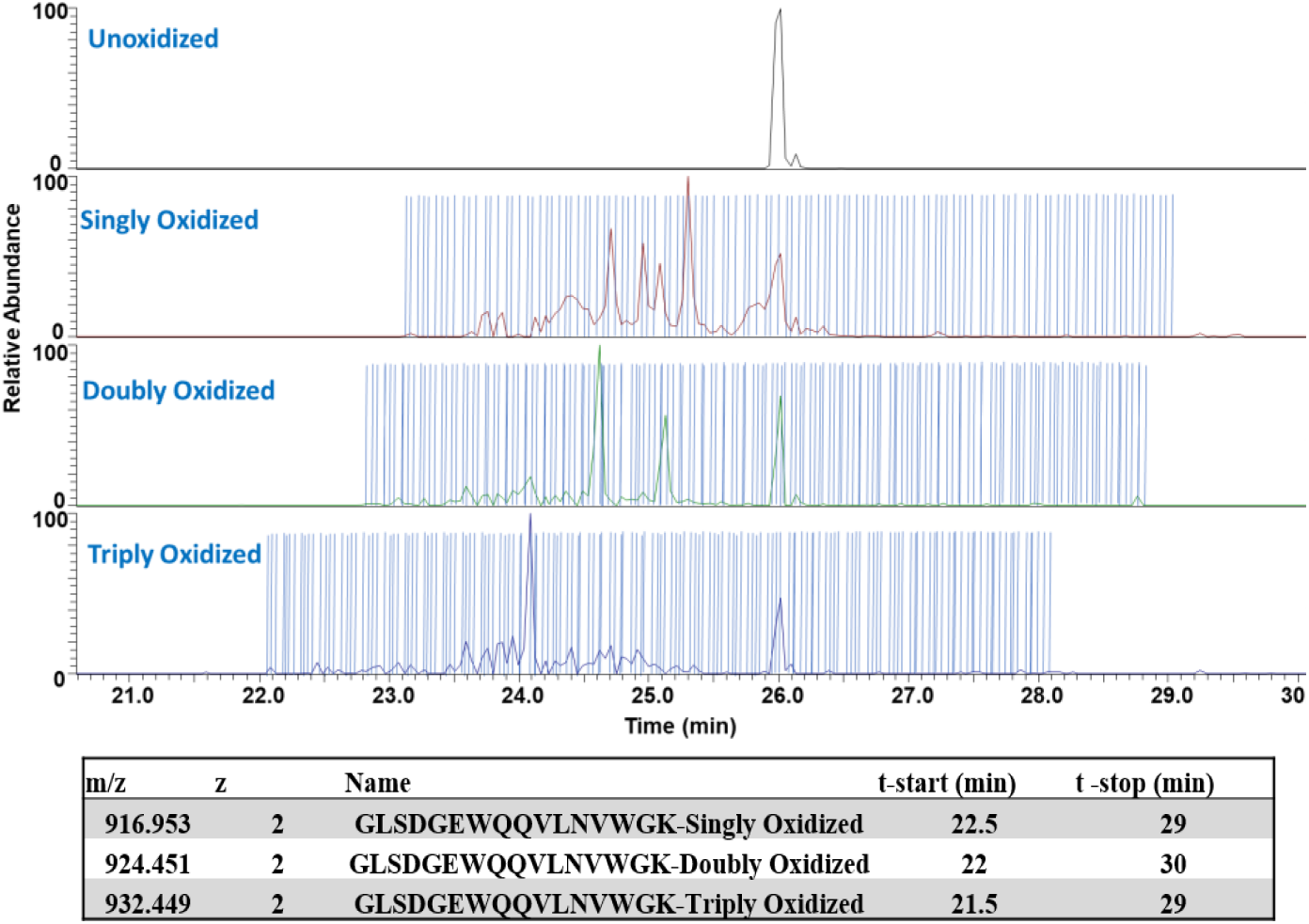
Representation of making mass inclusion list of a myoglobin peptide for a C18 RP ETD custom method. Each blue line represents an ETD MS/MS scan. This process has to be done for all the peptides in all of their oxidation states for each sample. It takes several hours to create a custom method with a full targeted mass inclusion list for each replicate.

If the peptide oxidation isomers can be made to perfectly co-elute chromatographically, while still retaining some separation ability for different peptides, these oxidation isomers can be accurately relatively quantified by ETD MS/MS product ion abundance from a single ETD spectrum acquired at any point during the elution of the peptide oxidation isomers, making the use of a standard data-dependent acquisition (DDA) method possible. Size exclusion chromatography (SEC) provides some low resolution separation of peptides, but allows peptide oxidation isomers to perfectly co-elute and be accurately quantified by DDA ETD MS/MS. However, SEC gives no on-column concentration, resulting in injection volume-dependent chromatographic resolution and poor sensitivity^33^. Additionally, difficulties exist in enacting robust SEC at low column IDs due, in part, to an amplification of the injection volume issue, further decreasing sensitivity and harming reproducibility and robustness.

Here, we developed a zwitterionic hydrophilic interaction chromatography (ZIC-HILIC) method to ideally co-elute peptide oxidation isomers while separating different peptides. This method only requires the use of 20-35 pmoles of sample using a 0.3 mm ID capillary column, with sensitivity rivaling that of standard nanoRPLC-MS using a 0.075 mm ID column. As ZIC-HILIC allows for on-column concentration, separation performance is largely independent of injection volume. This allows us to relatively quantify peptide oxidation isomers from any HRPF sample with a generic DDA ETD MS/MS method, using an ETD MS/MS spectrum acquired at any point across the single peptide oxidation isomer peak. This technology also greatly simplifies data analysis, both for manual analysis and future automation, making accurate HR-HRPF more approachable.

## Experimental Section

### Materials

Isomeric oxidized peptides (R(Hyp)MFAIWK, RPMYAIWK, and RPMFSIWK) were synthesized by GenScript (Piscataway, NJ), with purification and quantification performed by offline HPLC as previously described^32^. Bovine Serum Albumin (BSA) tryptic digest was obtained from Thermo Fisher Scientific (Waltham, MA). Myoglobin, catalase, trifluoroacetic acid (TFA), and sodium phosphate were purchased from Sigma-Aldrich (St. Louis, MO). Hydrogen peroxide (30%) was obtained from J.T. Baker (Phillipsburg, NJ). ZipTip with 0.6 µL C18 resin was purchased from Millipore sigma (Burlington, MA). Dithiothreitol (DTT), adenine and LC/MS-grade acetonitrile and water were obtained from Fisher Scientific (Fair Lawn, NJ). Sequencing grade modified trypsin was purchased from Promega (Madison, WI).

### FPOP

Myoglobin FPOP was performed as described previously^1^. Briefly, myoglobin samples were mixed to a final concentration of 10 µM with 17 mM glutamine, 1 mM adenine, 50 mM sodium phosphate buffer (pH 7.1), and 100 mM hydrogen peroxide immediately prior laser exposure. Each sample was pushed into a 100 μm i.d., 365μm o.d. fused silica capillary at 12.6 µL/min flow rate and exposed with COMPex Pro 102 KrF excimer laser (Coherent Inc., Santa Clara, CA). The laser was focused with a convex lens (Edmund Optics, Barrington, NJ) to a fluence of ~8.75 mJ/mm^2^. A 1 mm aperture (Edmund Optics, Barrington, NJ) was placed immediately prior the capillary to maintain a constant illumination window width, and the flow rate and illuminated length of capillary were adjusted for a 15% exclusion volume to ensure mostly single illumination of each sample volume while maintaining a high level of labeling^11,34^. Adenine radical dosimetry was conducted by using a Pioneer inline radical dosimeter (GenNext Technologies)^35^, and all sample dosimetry met the target value ΔAbs_265_ = 7.38 ± 0.285. After laser exposure and dosimetry, samples were quenched in 50nM catalase tetramer to enzymatically remove excess hydrogen peroxide and 20 mM methionine amide to scavenge secondary oxidants^36^.

### Sample Preparation for LC-MS

After performing FPOP on myoglobin samples, 50 mM Tris (pH 8.0) and 1 mM CaCl2 were added to the samples. The samples were denatured with heat at 95º C for 30 minutes in the presence of 5 mM DTT (unnecessary for myoglobin, but included to validate method performance in the presence of DTT). Then, the samples were immediately cooled down on ice for one minute. Trypsin was added at 1/20 enzyme/protein weight ratio to each samples following by 14 hours digestion at 37º C. The digestion was stopped by heating the samples at 95º C for 5 minutes. Digested samples were desalted using C18 ZipTips by first washing the tips 5 times with 10 µL elution buffer (95% acetonitrile + 0.05% TFA) each time.

The tips were equilibrated 5 times with 10 µL wash solution (5% acetonitrile + 0.05% TFA). Then, 10 µl of digested myoglobin samples were pulled from the samples. The tip was washed once with wash solution to remove excess salt. Bound peptides were eluted twice with 10 µL of elution solution each and immediately queued for ZIC-HILIC-MS/MS analysis.

### ZIC-HILIC-MS/MS

Freshly prepared samples were loaded on a SeQuant ZIC-HILIC capillary column (0.3 mm × 150 mm, 3.5 μm particle size, 200 A pore size, Millipore) using the capillary pump of a Dionex Ultimate 3000 Nano LC system with autosampler (Dionex, Sunnyvale, CA). Samples were loaded at 95% buffer B (acetonitrile, 0.05% TFA), 5% buffer A (water, 0.05% TFA) for 12 minutes at a flow rate of 2 µl/min. Initial elution then started by increasing the flow rate to 5 µl/min and moving to 20% A over a 4-minute linear gradient. Then, a linear gradient of 20% A to 40% A over 20 minutes was applied for higher resolution elution of most peptides. After elution, the wash gradient started by increasing to 60% A over 5 minutes and held isocratic at 60% A for 5 minutes. Buffer B was increased to 95% over 1 minute and held isocratic for 8 minutes to re-equilibrate the column for the next run.

Peptides were eluted directly into the HESI source of a Thermo Orbitrap Fusion (Thermo Fisher Scientific, Waltham, MA) instrument using a low flow rate emitter. The data was collected in positive ion mode with spray voltage at 3800 V in static mode. The ion transfer tube temperature was set to 300º C and vaporizer temperature was set at 30º C. For MS/MS scans, ETD was performed on +3 and higher charge state peptide ions with 100 ms ETD reaction time and 2.0e5 ETD reagent target. The maximum ETD reagent injection time was 200 ms. Product ions were detected using the orbitrap with resolution set at 60000 and AGC target at 2.0e5. For +2 charge state peptide ions, ETD with 20% EThcD supplemental activation (SA) energy was performed to promote dissociation of the charge-reduced radical cation. The reaction time, reagent targets and AGC targets were same as +3 and higher charge ETD method. Under these LC and ionization conditions, the 0.05% TFA was not found to cause issues with electrospray stability or source contamination after several extended series of runs. The MS method for evaluating sequence coverage of BSA samples was DDA with dynamic exclusion which excluded after 5 scans within a 30 sec window. The exclusion duration was 60 sec. For isomeric oxidized peptides, MS/MS was obtained using a targeted mass inclusion list to compare MS/MS spectra at different elution times. For myoglobin and BSA samples, DDA ETD MS/MS scans were done every 6 seconds on the peptide ions. A mass inclusion list that only covered expected oxidized versions of myoglobin peptides was used.

### C18 RPLC-MS/MS

C18 LC-MS/MS used an Acclaim PepMap 100 C18 nanocolumn (0.075 mm × 150 mm, 2 μm particle size, 100Å pore size, Thermo Fisher Scientific) coupled to a 300 μm i.d. ×5 mm C18 PepMap 100 trap column with 5 μm particle size (Thermo Fisher Scientific) to desalt and concentrate the samples before loading onto the C18 nanocolumn for separation. We used the capillary pump to load the samples onto the C18 trap column using buffer A (water + 0.05% TFA) and buffer B (acetonitrile + 0.05% TFA). We also used nanopump for chromatographic separation using mobile phase C (water+ 0.1% formic acid) and mobile phase D (acetonitrile+ 0.1% formic acid). First, the samples were loaded onto the C18 trap column in 2% B at 5 µl/min for 6 minutes. The trap column was then switched inline with the nanocolumn and trapped peptides were back-eluted onto the nanocolumn using the nanopump. Elution started by increasing solvent D in a linear gradient from 2% to 40% over 22 minutes. The gradient then ramped up to 95% D over 5 minutes and held isocratic for 3 minutes to wash the column. Buffer D was then decreased to 2% over 1 minute and held isocratic for 6 minutes to re-equilibrate the column for the next run. The samples were eluted directly into a nanospray source, where the spray voltage was set at 2400 V and ion transfer tube temperature at 300º C. Full MS scan was obtained from 150 to 2000 *m/z*. CID and ETD was performed on the 10 most abundant precursor ions of +2 charge and greater for peptide identification and sequence coverage analysis. For ions with +2 charge state, ETD was performed with 20% EThcD SA collision energy. The orbitrap resolution for both ETD and EThcD was 30000 with AGC target at 5e4 and maximum injection time of 100 ms.

### Data Analysis

Data analysis was performed as described previously^32,37^. Byonic version v2.10.5 (Protein Metrics) was used to identify BSA and myoglobin peptide sequences using a sequence database for each protein. For all myoglobin peptides detected, the major oxidation products detected were net additions of one or more oxygen atoms. In order to calculate average oxidation events per peptide, the area under the curve for peaks of unoxidized and oxidized peptides was used according to Eq. (1). In short, the oxidation events per peptide were calculated by summing the intensity of each peptide oxidation product multiplied by the number of oxidation events on the peptide required to generate that product and divided by the sum of I for all oxidized and unoxidized versions of that peptide, as shown in Equation 1. *P* represents the average oxidation events per peptide, and *I* is the area under the curve for peaks of oxidized and unoxidized peptides.

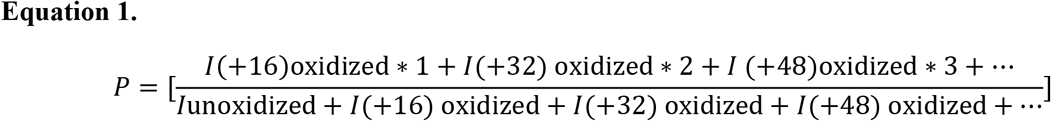

The amount of oxidation at residue level quantitation in a peptide was determined by the fragment ion (z or c ion) intensities of the peptide ETD fragmentation. The oxidation fraction of a given z or c ion was calculated by dividing the oxidized sequence ion intensity to the sum of the intensity of the corresponding oxidized and unoxidized sequence ion in a particular oxidized peptide. The relative oxidation fraction of each production *f*(*Ci*) was calculated using Equation 2 where *I*(*Ci*) is the intensity the designated product ion, either summed across all spectra for RPLC, or taken at any individual point for ZIC-HILIC.

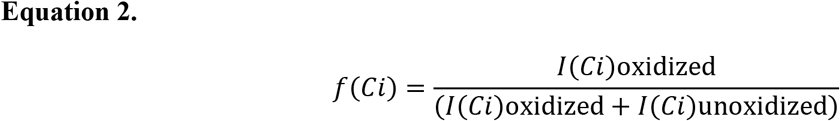

The absolute amount of oxidation of a given amino acid was determined by multiplying the average oxidation event of peptide by the absolute fractional oxidation of the corresponding sequence ions. As shown in Equation 3, *P* is the average oxidation event per peptide calculated from Equation 1, and the term in bracket is the fractional difference of two adjacent sequence ions, *f*(*C*_*i*_) and *f*(*C*_*i-1*_). In cases where ETD fragmentation ions are not adjacent in sequence, fractional oxidation for multiple contiguous residues within the peptide can was calculated by using non-adjacent ETD fragments in Equation 3.

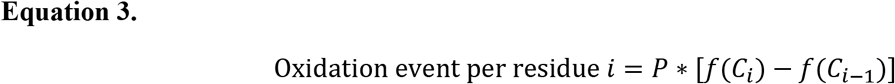

## Results and Discussion

### Co-elution and quantification of synthetic peptide oxidation isomers using ZIC-HILIC

We tested the co-elution of isomeric oxidized peptides by mixing three synthetic peptide oxidation isomers: R(Hyp)MFAIWK, RPMYAIWK, and RPMFSIWK. These isomers differ solely by the position of a single oxygen atom on each of three residues: Pro2, Phe4, or Ala5. Each peptide isomer was prepared at 10 µM and then mixed to a 1:1:1 molar ratio in 95% acetonitrile, meaning to the final concentration of each peptide was 3.33 µM in the mixture. Results from analysis by ZIC-HILIC and C18 RPLC were compared. As shown in Figure 2A, all three isomers were co-eluted in one peak using ZIC-HILIC gradient, while in Figure 2B, they separated into their three isomeric components by C18 RPLC as previously reported^33^. No separation can be directly observed by the peak shape in Figure 2A; the peak shape looks identical to that of any single component (data not shown).

**Figure 2.**
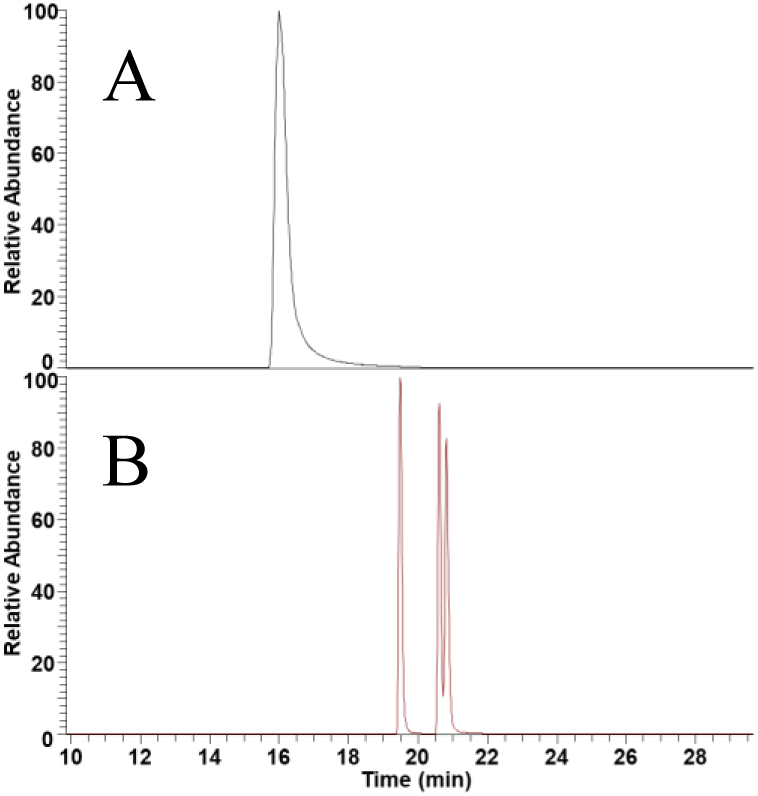
Chromatographic separation of synthetic isomeric peptide oxidation products. **A.** Capillary ZIC-HILIC yields co-elution of all three isomeric oxidized peptides. **B.** Isomeric oxidized peptides are separated by nano-C18 RP.

If the co-elution of all oxidation isomers is ideal in ZIC-HILIC as it appears in Figure 2A, accurate quantification of each isomer should be achievable by ETD MS/MS at any point in the chromatogram, within the limits of sensitivity. However, if co-elution of oxidation isomers is not ideal, systematic skews in quantification will be detected^33^. Figure 3 shows a selected product ion chromatogram, comparing the elution profile of oxidized peptide precursors that produce an oxidized c2 ion (indicating oxidation on the P) with the elution profile of oxidized peptide precursors that produce an unoxidized c2 ion (indicating oxidation on the F or A). The two traces are indistinguishable, further supporting the conclusion that the oxidation isomers are co-eluting ideally. ETD MS/MS spectra were obtained at five elution times spanning the peak: 35% and 70% peak height at leading edge, 100% peak height, and 35% and 70% peak height at tailing edge. The fractional oxidation of each amino acid residue in the mixture of oxidation isomers was measured based on relative abundances of the oxidized vs. unoxidized product ions; one such spectrum is shown in Supplementary Information, **Figure S1** for illustrative purposes. As shown in Figure 3, all time points generated quantification of each oxidation isomer that was very close to the theoretical value.

**Figure 3.**
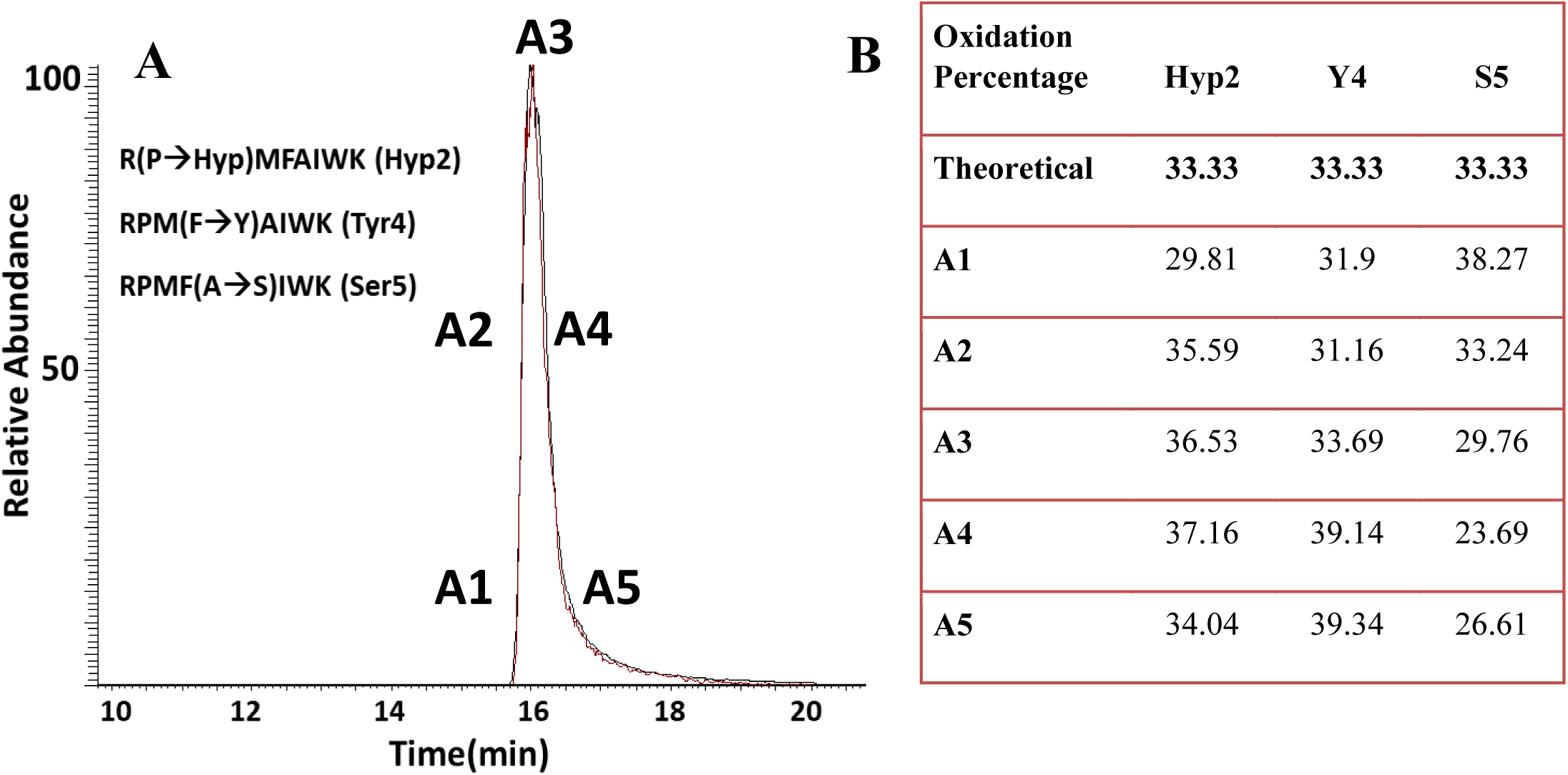
Measured oxidation of synthetic peptide isomer set RPMFAIWK, calculated using ETD spectra at various ZIC-HILIC retention times. **(A)** Extracted product ion chromatogram of c2 product ion from oxidized peptide precursor. Black trace shows the unoxidized c2 ion, specific for the two isomers containing oxygen on F or A, while the red trace shows the oxidized c2 ion, specific for oxidation of P. Two traces are overlapping, indicating co-elution of the oxidation isomers. Markers on the peak indicate ETD spectra used for quantification. **(B)** Quantification of each oxidation isomer at each of five different elution times, using ETD product ion abundances.

In order to test the robustness of ZIC-HILIC ETD method, we mixed the peptide isomers in six different relative ratios of each isomer, while maintaining a constant total peptide concentration: 1:3:1, 1:1:3, 3:1:1, 3:1:2, 1:4:1, and 1;1:1 of Hyp2, Y4, and S5, respectively. We then measured their oxidation percentages from single ETD spectra at five different retention times across the elution profile, and compared the theoretical and experimental quantitation, shown in Figure 4. The measured oxidation fraction corresponds very strongly to the theoretical oxidized fraction, demonstrating ideal co-elution of these oxidation isomers in an adsorptive capillary LC format, using sample amounts comparable to those used for RPLC. No systematic error in quantification based on peptide ratio or the time at which the ETD spectrum was taken is apparent.

**Figure 4.**
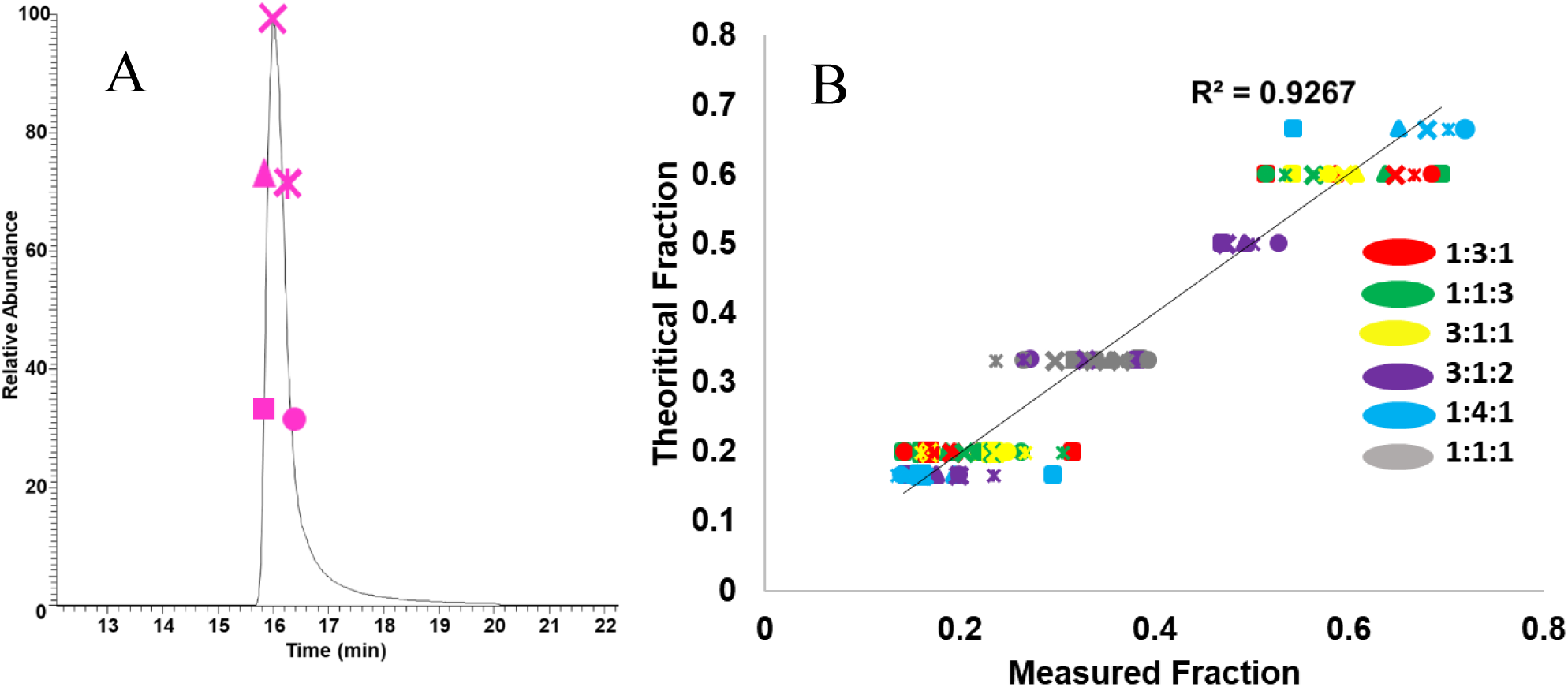
Measured oxidation of RPMFAIWK oxidation isomers at varying relative concentrations calculated using ZIC-HILIC ETD spectra at five different retention times. **(A)** Representative ZIC-HILIC selected ion chromatogram of the peptide oxidation isomers. Each shape represents a different retention time sampled by ETD for quantitation. **(B)** Measurement of oxidation of three peptide oxidation isomers mixed in six different relative ratios. The x-axis represents the oxidation of each amino acid as measured by ZIC-HILIC ETD, while the y-axis represents the theoretical oxidation of that amino acid in the synthetic mixture. The diagonal line represents x=y. Different marker shapes represent different times for ETD MS/MS spectrum acquisition as shown in **A**; different colors represents different ratios of synthetic oxidation products listed as Hyp2:Tyr4:Ser5. No systematic error is apparent; the measurements all cluster around the theoretical value.

### Ability of ZIC-HILIC to Separate Non-Isomeric Peptides for High Sequence Coverage

After achieving the co-elution of isomeric peptide, we tested the ability of ZIC-HILIC in separating different peptides using both a commercial BSA tryptic digest standard and myoglobin tryptic digest peptides. As shown in Figure 5A and 5B, separation of non-isomeric BSA and myoglobin peptides was achieved by ZIC-HILIC. The sequence coverage of commercial BSA tryptic peptide standard achieved by ZIC-HILIC as shown in Supplementary Information **Figure S2** was 81.06%, compared to 76% with C18 RP LC-MS/MS using the same amount of injected sample (200 fmoles). We also tested the separation and sequence coverage with an in-house tryptic digestion of myoglobin. For myoglobin ZIC-HILIC sample preparation, C18 solid phase extraction with gentle washing conditions and high organic elution (95% acetonitrile+0.05% TFA) was used in order to desalt 10 µL of the tryptic digest, preventing a salt phase partitioning problem in HILIC-compatible high organic solvent while maintaining sensitivity. As shown in **Figure S3**, the sequence coverage of myoglobin achieved by ZIC-HILIC was 98.70%, compared to 95.45% with C18 RP LC-MS/MS. Capillary ZIC-HILIC using the gradient described here was able to generate sequence coverages comparable to nano-C18 RP LC-MS/MS. In order to ensure the sample preparation by C18 solid phase extraction does not affect the co-elution and accurate quantification of isomeric oxidized peptides, we used this solid phase extraction-ZIC-HILIC protocol on our synthetic mixture of isomeric peptide oxidation products. The peptides were mixed in 1:1:1 molar ratio in aqueous solution and subjected to C18 solid phase extraction as described. As shown in **Figure S4**, we obtained perfect co-elution and accurate quantification of all peptide oxidation isomers.

**Figure 5.**
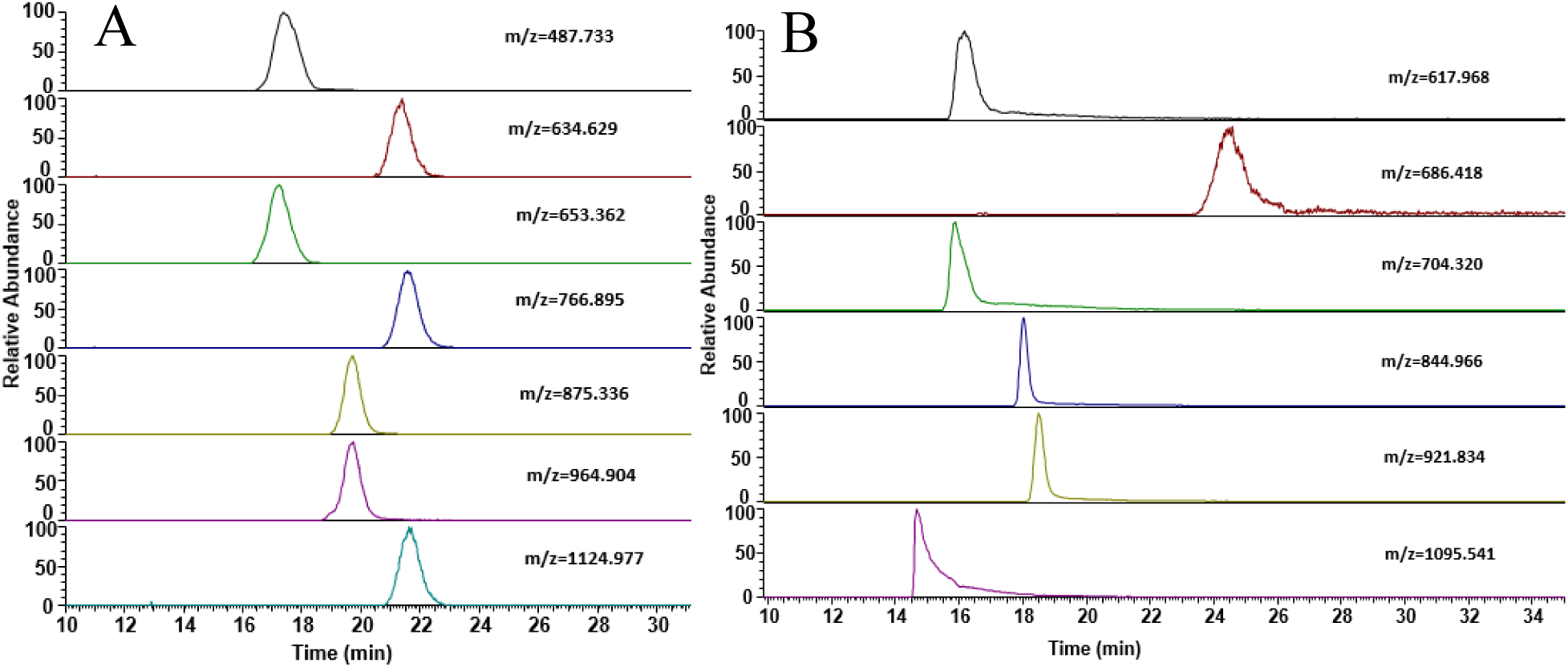
Separation of non-isomeric tryptic peptides. Selected ion chromatograms for various peptides are shown for ZIC-HILIC separation of tryptic digests of **(A)** BSA and **(B)** myoglobin.

### ZIC-HILIC for High-Resolution HRPF of Myoglobin

In order to test the general applicability of ZIC-HILIC ETD method, FPOP was performed on myoglobin as described above, and the oxidized peptides were analyzed by both ZIC-HILIC ETD LC-MS/MS and C18 RP ETD LC-MS/MS. Figure 6 demonstrates the methodological improvement in ZIC-HILIC ETD HR-HRPF as compared to traditional C18 RPLC-based methods. As shown in Figure 6A, all FPOP +16 Da oxidation isomers of the VEADIAGHGQEVLIR peptide and c10 oxidized and unoxidized ions are co-eluting ideally off of the ZIC-HILIC column, allowing us to quantify accurately by acquiring an ETD spectrum at any point during the elution time, which can be easily achieved by data-dependent acquisition (with optional targeted inclusion lists to improve depth of coverage for expected oxidation products). By comparison, C18 RPLC as shown in Figure 6B separates oxidation isomers and the oxidized and unoxidized ions, complicating data acquisition and analysis.

**Figure 6.**
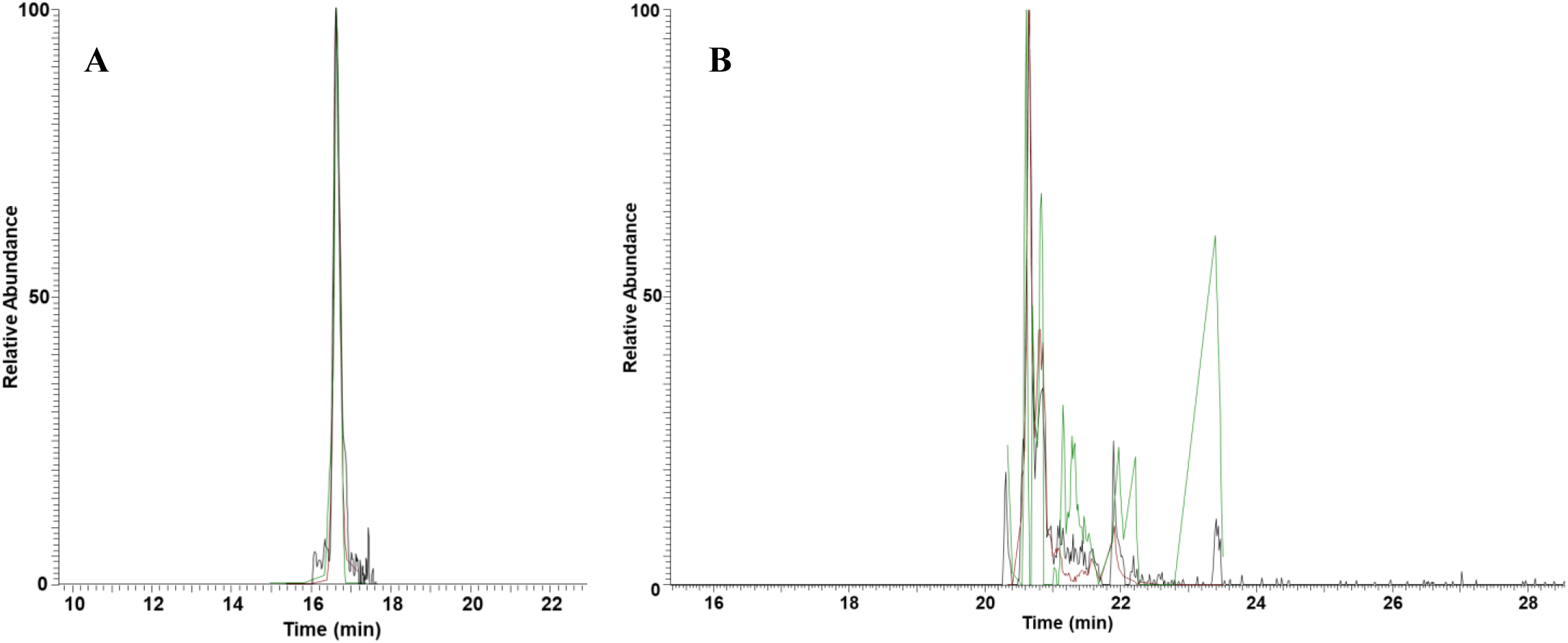
Extracted product ion chromatogram of c10 product ion for myoglobin peptide (VEADIAGHGQEVLIR) M+16 isomers via ZIC-HILIC and C18 RPLC. Black trace shows selected ion chromatogram of singly oxidized VEADIAGHGQEVLIR isomer peptides. Red trace shows the unoxidized c10 ion, while the green trace shows the oxidized c10 ion. **(A)** In ZIC-HILIC chromatography, the three traces are overlapping, indicating co-elution of peptide oxidation isomers. Accurate quantification of oxidation at the residue level could be obtained by a data-dependent ETD MS/MS spectrum obtained at any point along the single peak. **(B)** In C18 RPLC, there is a significant spread in the elution times of various peptide oxidation isomers. Accurate quantification of these isomers would require scheduling of ETD MS/MS across an almost four minute elution time window.

As shown in Figure 7, ZIC-HILIC LC-MS of FPOP-oxidized myoglobin yields HRPF peptide-level quantification that is identical to that of traditional C18 RPLC-MS. We observed 10 peptides, and we obtained similar quantitative HRPF results using both methods. Student’s t-test was run between oxidation event of each peptide obtained using each method and no statistically significant differences were observed (α=0.05).

**Figure 7.**
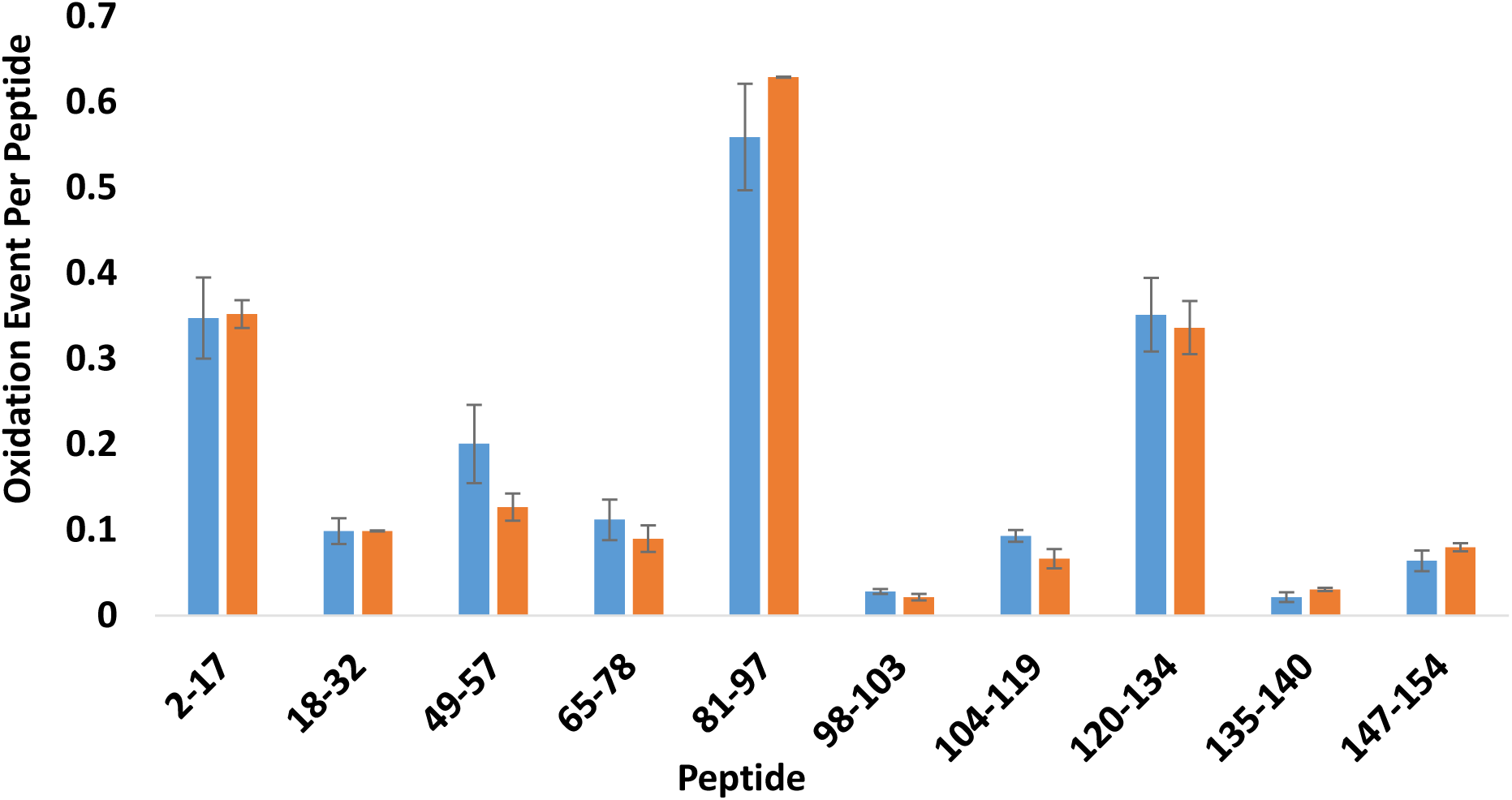
Peptide level comparison of Myoglobin peptides using ZIC-HILIC (blue) VS. C18 chromatography (orange). Both methods generated statistically indistinguishable quantification for each peptide (α = 0.05).

We tested the ability of ZIC-HILIC ETD LC-MS/MS to quantify the oxidation of each amino acid in myoglobin using a simplified DDA ETD MS/MS approach, using a mass inclusion list that only covers the theoretical oxidized versions of myoglobin peptides to improve depth of coverage. As shown in Figure 8, we could accurately quantify the average oxidation events across 22 segments covering 9 out of 10 detected peptides using ZIC-HILIC ETD LC-MS/MS. Most of these segments represent single amino acids, allowing for true HR-HRPF analysis using a fairly generic data-dependent acquisition method in a capillary LC format that allows for on-column sample concentration and sensitivity comparable to traditional C18 nanoRPLC-MS.

**Figure 8.**
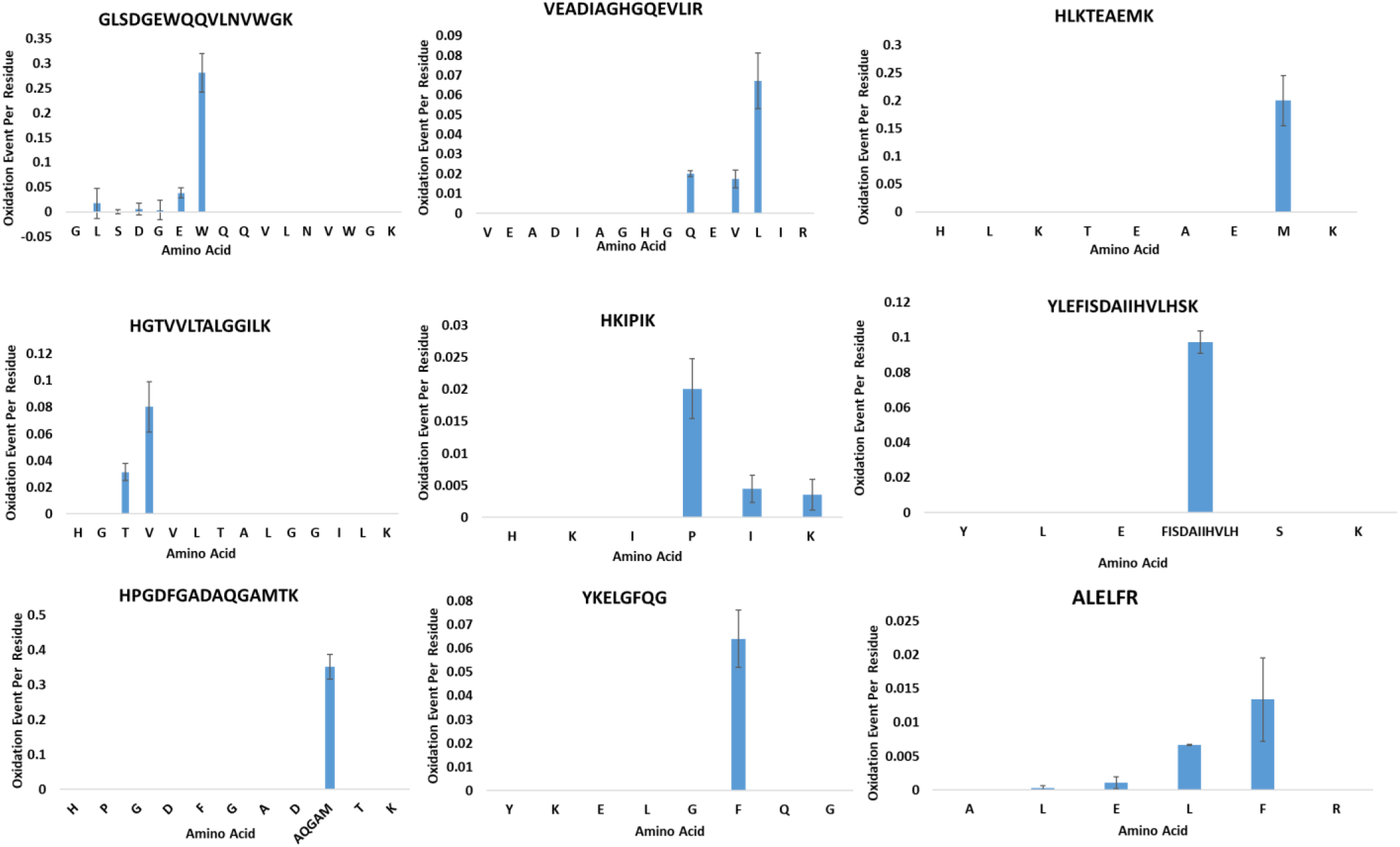
Amino acid level analysis of myoglobin peptides using ZIC-HILIC.

We compared our oxidized result with the previously reported HR-HRPF results of myoglobin conducted using C18 RPLC, albeit with a much more thorough multi-point FPOP protocol^3^. Most of the oxidized residues using ZIC-HILIC method was found oxidized in the previously reported result. As shown in **Figure S5**, the oxidation event per residue of ZIC-HILIC is correlated well with the radical dose response of the same residues from the previous report (R^2^ = 0.7375), even though the data generated here were taken at a single radical dose, and the ZIC-HILIC results could be obtained and analyzed in a small fraction of the time required for C18 RPLC-MS/MS.

## Conclusion

It was demonstrated in this work that our ZIC-HILIC ETD method provides ideal co-elution of peptide modification isomers, allowing for accurate quantification of oxidation at the residue level using a single ETD MS/MS spectrum acquired anywhere across the peak. In traditional C18 RPLC the peptide modification isomers do not co-elute. Therefore, ETD spectra must be scheduled manually to properly sample all oxidation isomer peaks eluting across the entire elution window. Improper sampling results in erroneous quantitation of oxidation isomers. This requires multiple runs of each HR-HRPF sample: one run to identify the detected oxidation isomers and their elution times, and then one or more runs with a custom method to perform scheduled ETD of each isomer at a sufficient scan frequency to sample all peaks^3,33,38,39^. ZIC-HILIC reduces method development time significantly. Unlike previously published SEC-ETD MS/MS^33^, it is also feasible with HRPF experiments when the protein quantity is limited due to the ability to perform capillary-scale LC with on-column concentration. ZIC-HILIC requires only a few pmoles of sample and provides similar sequence coverage and oxidation quantification compared to the C18 RP method.

The other benefit of our developed ZIC-HILIC method is the significant reduction in HRPF data processing time. In C18 RPLC, the *m/z* and *z* must be confirmed for each eluting peak to prevent misidentification of isobaric analytes eluting within the broad window in which peptide oxidation products are commonly found as a peptide oxidation product. This is especially challenging for oxidation products of low abundance, as they sometimes do not exhibit strong isotopologue signals resulting in insufficient information for confident charge state assignment. Using ZIC-HILIC ETD MS/MS to quantify the unoxidized and isomeric oxidized products at the peptide level, only one peak *m/z* and *z* confirmation is required for each group of oxidized isomer products due to the co-elution of oxidized isomers in one single peak. In our hands, data from the C18 RPLC method takes about three times as long to analyze as our newly developed ZIC-HILIC method. This large difference in the time and complexity of data analysis with the standard RPLC method, compared to our new ZIC-HILIC method, has historically been a hindrance on the adoption of HRPF-FPOP in more laboratories. The use of the ZIC-HILIC method should entice the adoption of this method by more of the scientific community and should allow for the development of automated peak-picking and analysis software which would make this method even more widely desirable.

ZIC-HILIC analysis of FPOP data may also be useful for simpler automated oxidation quantification at the peptide level. Automated peak picking of C18 RPLC-MS oxidation isomers is very challenging due to the complex and partially resolved nature of oxidation isomers as shown in Figure 5B, resulting in peaks that rarely show the ideal peak shape in the selected ion chromatogram. ZIC-HILIC LC-MS elutes all oxidation isomers as a single peak that is relatively well-shaped, making automated peak picking and integration much easier. This should result in better data analysis in much shorter periods of time, making it easier to study larger and more complex proteins as well as greater density which can be used for more in depth studies of these proteins and their binding partners. The improved peak shapes may also make peak identification and integration using automated tools easier and more accurate.

## Acknowledgments

This work was funded by the National Institute of General Medical Sciences (R01GM127267).

## Data Sharing

All LC-MS data sets used in this publication are available at https://massive.ucsd.edu, dataset identifier MSV000084605.

