## Supplementary Information for "Rapid Quantification of Peptide Oxidation Isomers from Complex Mixtures"

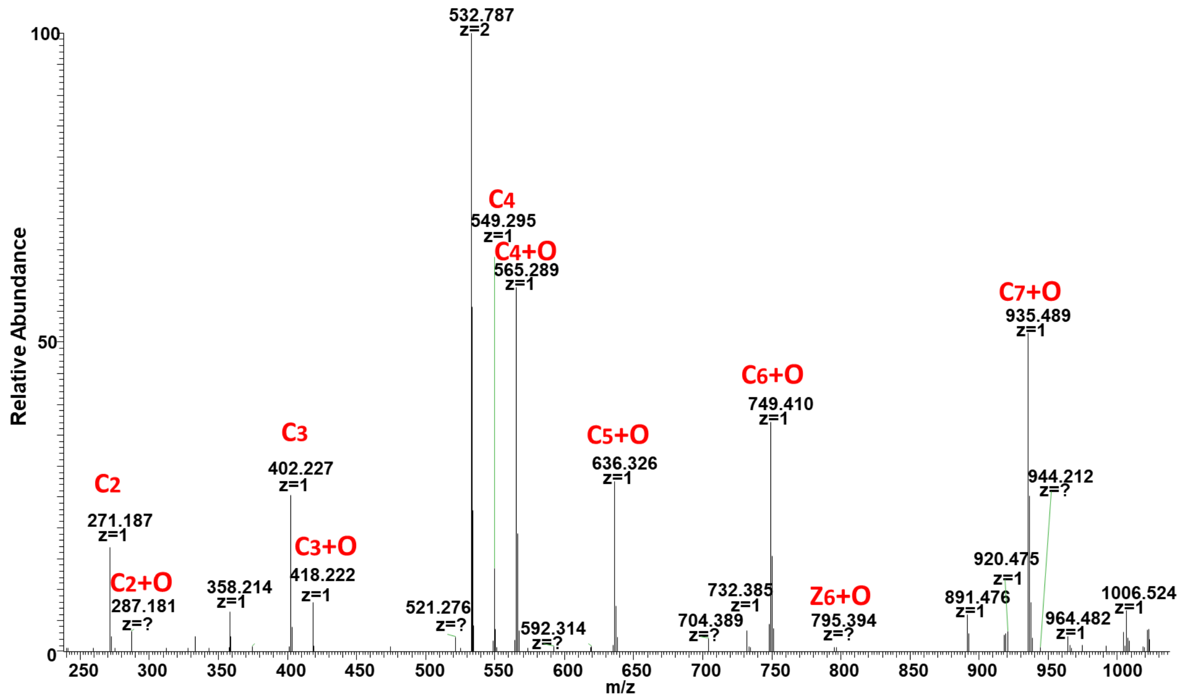


**Figure S1. Annotated ETD spectrum the 1:1:1 mixture of synthetic peptide isomers, obtained at maximum LC peak height.**

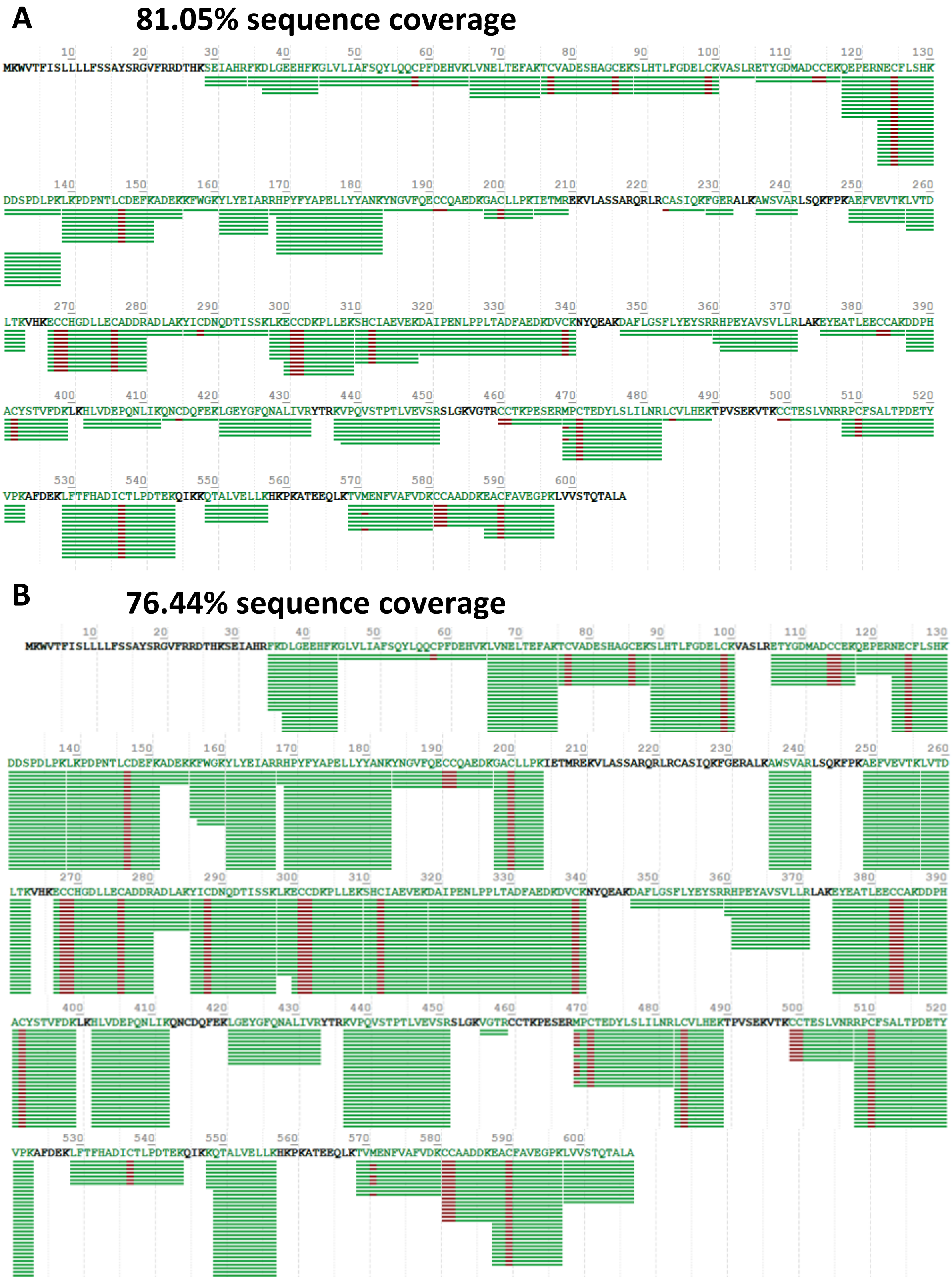


**Figure S2. Sequence coverage comparison of BSA between (A) ZIC-HILIC and (B) C18 RPLC.** While the repeat count is higher for C18 RPLC, the sequence coverage is higher for ZIC-HILIC.

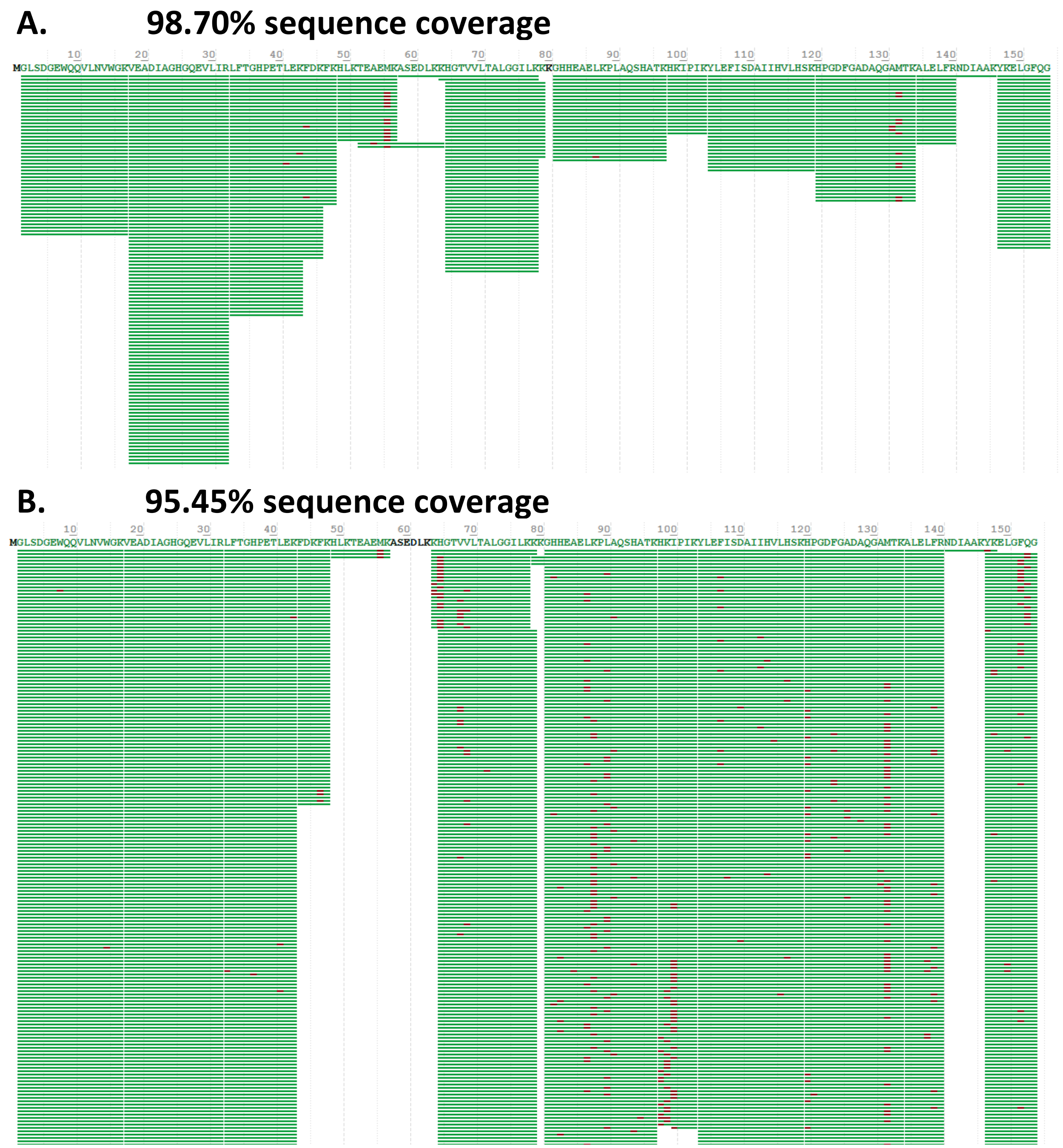


**Figure S3. Sequence coverage comparison of oxidized myoglobin between ZIC-HILIC and C18 RP.** (A) Myoglobin sequence coverage achieved by ZIC-HILIC. (B) myoglobin sequence coverage using C18 RP.

A.

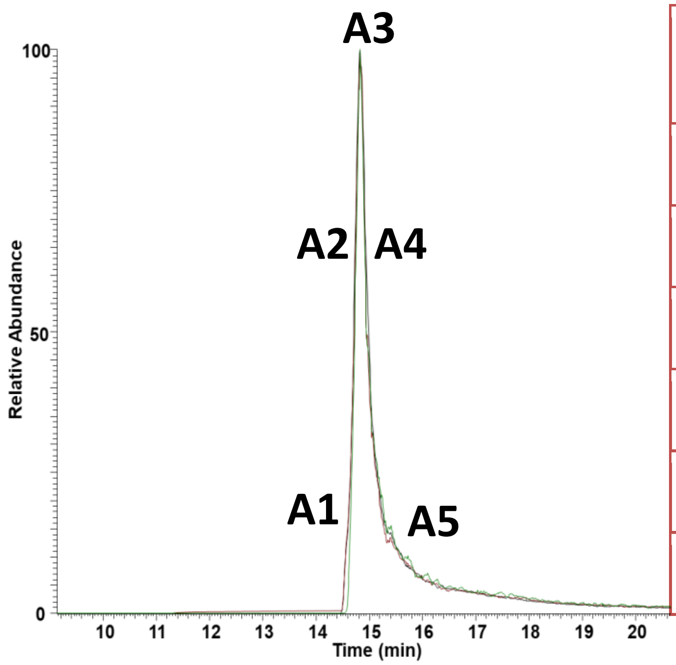


B.

| Oxidation Percentage | Hyp2 | Y4 | S5 |
|---|---|---|---|
| **Theoretical** | **33.33** | **33.33** | **33.33** |
| **A1** | 30.53 | 23.37 | 46.08 |
| **A2** | 26.92 | 36.86 | 36.20 |
| **A3** | 32.96 | 32.96 | 34.07 |
| **A4** | 31.25 | 43.96 | 24.78 |
| **A5** | 36.18 | 39.59 | 24.22 |

**Figure S4. Measured oxidation of synthetic peptide oxidation isomers after being prepared by solid phase C18 extraction. (A)** Extracted product ion chromatogram of c2 product ion from oxidized peptide precursor. Black trace shows selected ion chromatogram of oxidized isomer peptide. Red trace shows the unoxidized c2 ion, specific for the two isomers containing oxygen on F or A, while the green trace shows the oxidized c2 ion, specific for oxidation of P. Three traces are overlapping, indicating sample preparation using C18 solid phase extraction does not affect the co-elution of isomeric peptides. Markers on the peak indicate ETD spectra used for quantification. **(B)** Quantification of each oxidation isomer at each of five different elution times, using ETD product ion abundances. No systematic error is apparent, indicating solid phase C18 extractions using ziptip does not cause inaccurate oxidation measurements.

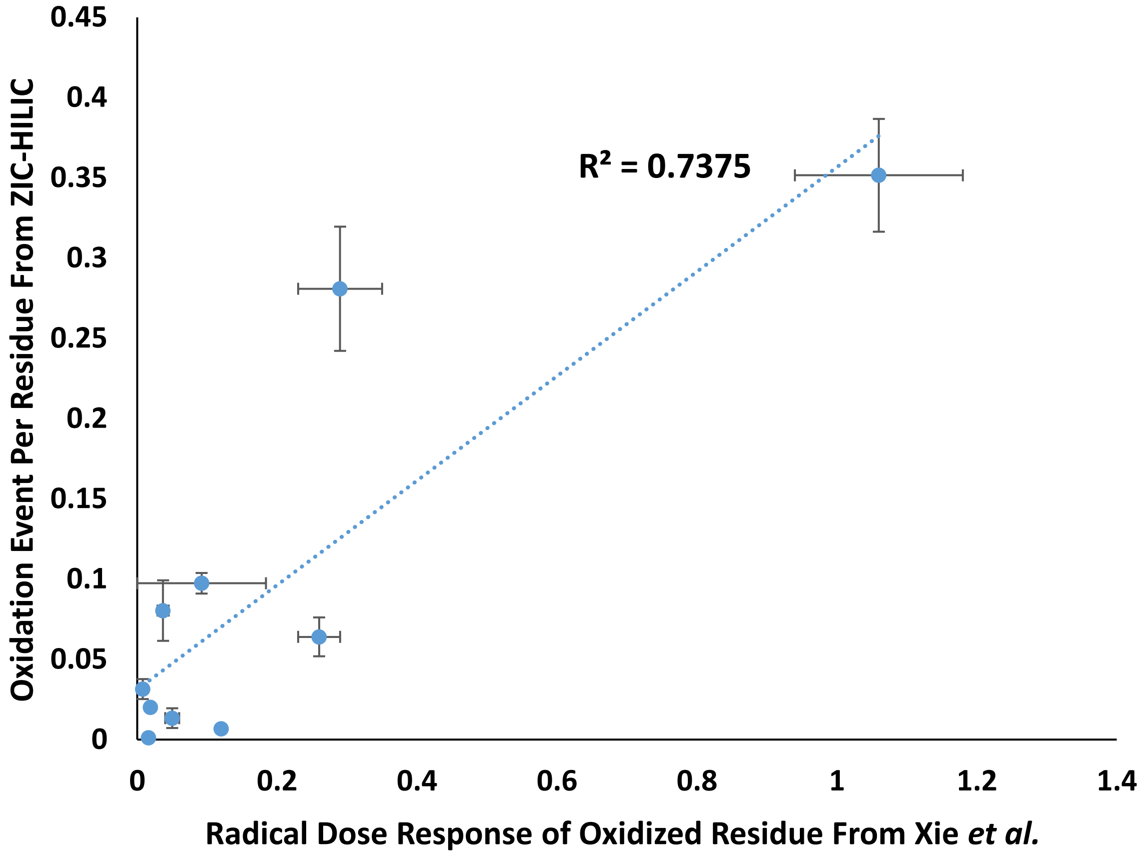


**Figure S5. Comparison of oxidation event per amino acid via ZIC-HILIC with radical dose response of them via previous method[1].**
